# A recurrent neural network model of chronic pain development and recovery

**DOI:** 10.64898/2026.04.18.719337

**Authors:** Katherine Huang, Guy Marmor, Tjitse van der Molen, Zongren Zhang, Paulina Gicqueau, Javier Reveles, Kareena Morrissey, Justin Tang, Lucy Lu, Kelly Ilmi, Jacob Lue, Gabriela Barba Zuniga, Michael B. Miller, Kenneth S. Kosik, Henry Yang, Tyler Santander, Francesco Bullo, Paul Hansma

## Abstract

Chronic pain presents a leading challenge in the world today for both clinicians and researchers. Because chronic pain is difficult to explain and treat, it is often managed with opioids despite providing limited relief and contributing to dependence and misuse. Persistent pain can be maintained by altered central nervous system processing even in the absence of distinct tissue damage or disease, which may limit the efficacy of conventional pharmacological therapies that target nociceptive signal transmission rather than maladaptive central nervous system dynamics often present in those with chronic pain. Although neuroimaging studies have identified this shift from nociceptive to emotional circuits during pain chronification, a quantitative framework linking these neural changes to longitudinal pain trajectories or recovery is lacking. We present a parsimonious firing-rate model that can account for the development of and recovery from chronic pain, which is based on the theoretical framework established by Wilson and Cowan. The model provides a quantitative explanation of how sensitization, anxiety, and fear maintain pain even after an injury has healed, and how calming stimulus downregulates these processes to facilitate recovery. A study applying the same principles as the model produced an average pain decrease of 3.5 on the Visual Analog Scale (VAS), with all subjects experiencing a reduction in pain. These results, coupled with our model and findings in prior studies, suggest that increasing calming stimulus can reduce pain without necessitating pharmacological or invasive, resource-intensive interventions.

## Introduction

The recent success of Pain Reprocessing Therapy (PRT) in reducing chronic pain (1–4) raises the question of how the relatively rapid reductions in long-term chronic pain can occur with non-pharmacological interventions. This could be explained in part by the presence of nociplastic pain, where altered neural processing drives persistent pain in the absence of tissue damage or disease(5–10). The model presented here explains how the maladaptive neuroplasticity of nociplastic pain can contribute to the chronification of pain even beyond the resolution of the initial injury(11). A tremendous amount has been learned about nociceptive and neuropathic pain through biochemical and neurotransmitter-oriented research focusing on molecular mechanisms such as inflammatory mediators(12), ion channels(13), synaptic modulators(14), and glial activation(15). However, additional research is needed to explain nociplastic pain, as nociceptive and neuropathic mechanisms do not account for the persistence of pain in the absence of ongoing peripheral input, nor do they explain why recovery is achieved with non-pharmacological interventions such as PRT. Development of chronic pain due to CNS changes is typically less responsive to standard drug-based interventions, as the absence of identifiable tissue damage leaves no clear therapeutic target(16). Thus, alternative non-pharmacological methods must be explored to address chronic pain for long-term recovery(17).

Here, we introduce a new quantitative model based on the pioneering work of Wilson and Cowan(18) for chronic pain that explains the chronification and persistence of pain due to neuroplastic changes and the recovery from pain with approaches such as PRT(2).

Specifically, we present a firing-rate recurrent neural network (FRNN) model of chronic pain that captures the temporal evolution of neural activity, providing a systems-level framework for the development and treatment of chronic pain. This systems-level framework offers a fundamental understanding of nociplastic pain mechanisms and identifies new research targets for developing more effective chronic pain treatment programs.

Additionally, we present results from chronic pain recovery studies designed using the same principles as our model framework, in which the primary intervention aimed to reduce the fear and anxiety associated with sensation. Almost all participants, with a variety of chronic pain conditions, experienced an overall reduction in pain.

## 2. Methods

### 2.1. A firing-rate recurrent neural network (FRNN) model for Chronic Pain

Here, we present a firing-rate recurrent neural network (FRNN) model of chronic pain that captures the temporal evolution of neural activity, providing a systems-level framework for the development and treatment of nociplastic pain (Fig 1). The two inputs in the mathematical model are pain stimulus (ps) and calming stimulus (cs). Pain stimulus is any input, external or internal, that triggers a pain pathway in the nervous system and is perceived as painful. Calming stimulus is any input, external or internal, that reduces pain-related sensitization, anxiety, or fear (SAF). Pain self-efficacy is an example of a calming stimulus(19).

**Fig 1.**
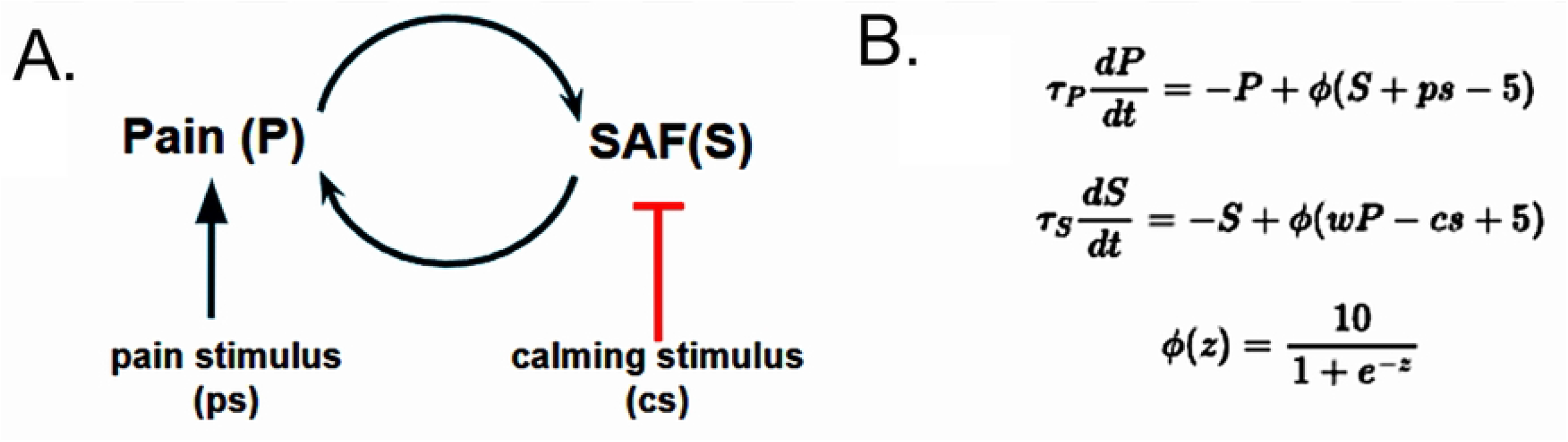
A firing-rate recurrent neural network model of nociplastic pain. **A**. Pain stimulus from nociceptor activation or neuropathy causes nociceptive or neuropathic pain through a connection (blue arrow) that involves gates and chemical systems that determine how much, if any, Pain (P) results from the pain stimulus. All nociceptive and neuropathic pain originates from pain stimulus. Sensitization, Anxiety, and Fear (SAF) facilitate the transition from acute to chronic pain and can maintain the Pain even in the absence of pain stimulus. Calming stimulus decreases Sensitization, Anxiety and Fear (SAF). The blue arrows are excitatory inputs. The red bar is an inhibitory input. **B**. Two-dimensional model equations. SAF is abbreviated as S in the equations. Additional parameters are defined in Section 2.2. This firing-rate model for chronic pain is based on the robust experimental and theoretical evidence that neural firing in the pain processing system correlates with pain (see below for references and details). The model is based on the framework of the Apkarian group’s neuroimaging research, which demonstrated that chronic pain development cannot be explained solely by pain stimulus(20). Altered neural firing, especially in regions associated with emotion, is largely involved in the transition from acute to chronic pain.

The firing-rate model characterizes the evolution of the firing rates of neurons associated with the experience of chronic pain and those associated with sensitization, anxiety, and fear. Specifically, we denote the average firing rate activations of these two populations as the Pain firing rate “P” and the SAF firing rate “S.” Accordingly, we propose an aggregate dynamical model describing the two neuronal populations as a two-dimensional dynamical system. This concept of an aggregated neural circuit is grounded in the classical neural circuit modeling framework established by Wilson and Cowan in 1972(18).

There is justification for all the connections in this model from published research:

1. The connection from **pain stimulus (ps) → Pain** describes pain generated by the activation of nociceptors in response to a particular stimulus or stimuli. A pain stimulus can be provoked directly or indirectly. Noxious stimuli, such as physical trauma, directly activate nociceptors to generate the experience of pain(21). Non-noxious stimuli, such as stress, indirectly activate nociceptors by eliciting physiological responses that cause the release of signaling molecules, which then can bind to and activate nociceptors to generate pain(22). In the context of our model, any stimulus that activates nociceptors to create the experience of pain is considered a pain stimulus.
2. The connection from **Pain → SAF** describes how persistent pain contributes to sensitization, anxiety, and fear. Prolonged nociceptor activation lowers firing thresholds and can cause normally innocuous stimuli to be perceived as painful(23), thus increasing sensitization[2,8,35,64]. Persistent pain may also cause emotional distress, increasing fear and anxiety(24). This relationship demonstrates the adverse effects of chronic pain on mental health, particularly anxiety(25).
3. The connection from **SAF → Pain** describes how increases in sensitization, anxiety, and fear further amplify pain, reinforcing a self-perpetuating cycle that contributes to the persistence of chronic pain. Anxiety(24,26–29) and pain-related fear(30–32) have been found to increase pain perception and reduce pain thresholds. Heightened sensitization can cause non-threatening stimuli to be misinterpreted as painful(33), further reinforcing fear and anxiety(32,34). Fear and anxiety also perpetuate one another through mechanisms demonstrated by the fear-avoidance model(35–37). The growing literature on the co-occurrence of chronic pain and anxiety supports our bidirectional model presented in connections 2 and 3(38–40).
4. The connection from **calming stimulus (cs) → SAF** demonstrates how sensitization, anxiety, and fear can be attenuated by downregulating the sympathetic nervous system and enhancing perceptions of safety. Physiologically calming stimuli, such as biofeedback, activate the parasympathetic nervous system to promote relaxation, thereby reducing cytokine production and inflammation associated with chronic pain (41–44). Cognitively calming stimuli, such as Pain Neuroscience Education (PNE), reduce pain-related fear and kinesiophobia by helping reframe misconceptions about pain (45–47). By modulating these drivers of chronic pain with calming stimuli, the experience of pain is decreased. A recent breakthrough study using intracranial implanted electrodes in people with chronic pain provides a compelling example of an internal calming stimulus(48). Participants reported pain metrics simultaneously with ambulatory neural recordings multiple times a day over months. Results indicated lower pain levels were associated with increased oscillatory power in the orbitofrontal cortex (OFC), a region implicated in processes related to affective regulation. Thus, in the context of our model, this increased oscillatory power would be a component of calming stimulus.

The SAF firing rate (S) includes not only sensitization, but also anxiety and fear, as all three amplify pain on distinct timescales. Anxiety and fear, in addition to contributing to sensitization over time, can increase pain within minutes, while sensitization, encompassing both peripheral and central sensitization, changes over longer time scales. If anxiety is reduced, pain is also reduced(43) (Fig 2).

**Fig 2.**
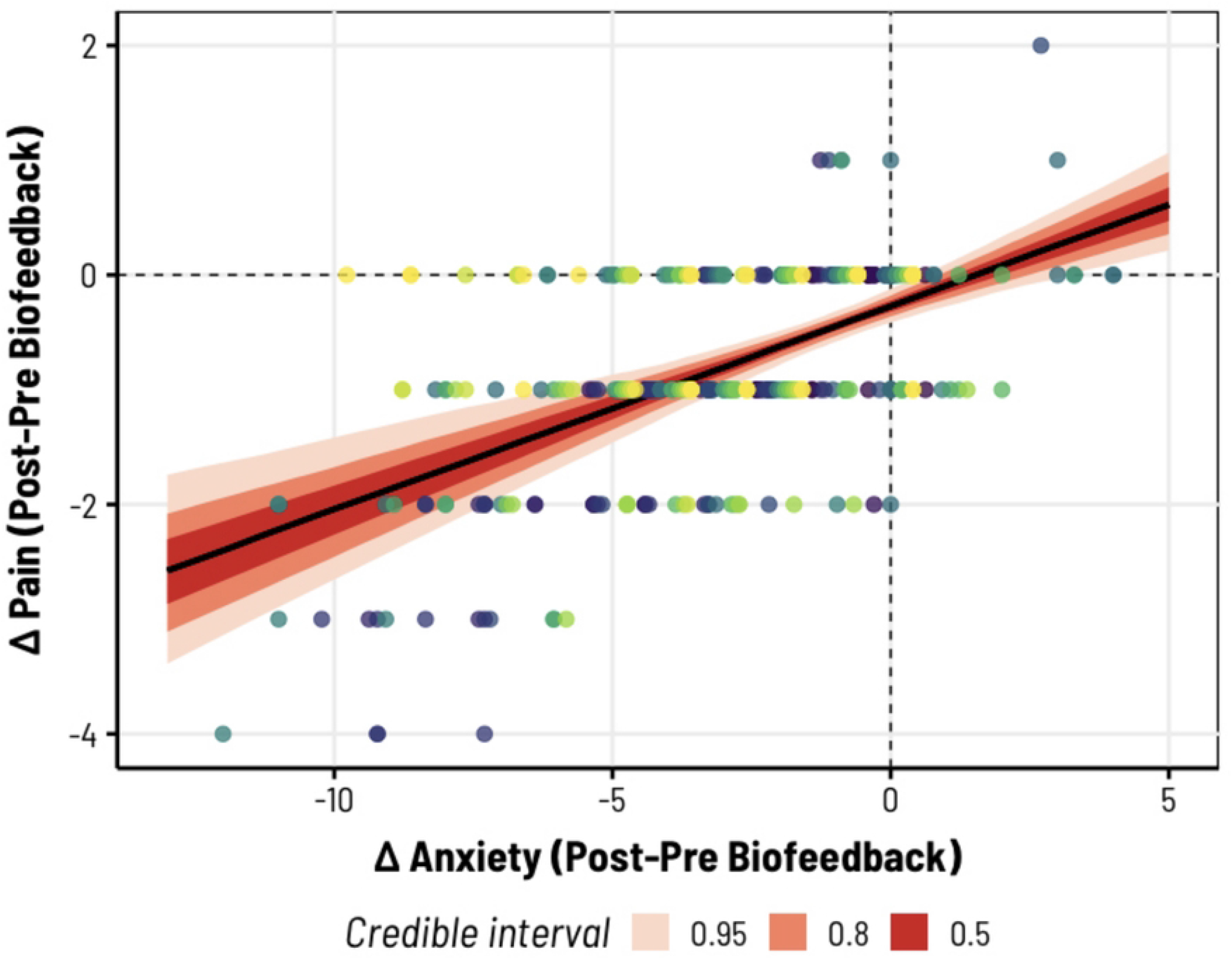
Results for changes in anxiety and pain from 10-minute biofeedback sessions. Decreases in pain (VAS) were correlated with decreased anxiety (SF-STAI) from 10-minute hand-temperature biofeedback sessions. Different colored data points correspond to different subjects. The population-level slope is displayed with Bayesian highest density intervals (shaded in red) (Data re-analyzed from “Home-Use and Portable Biofeedback Lowers Anxiety and Pain in Chronic Pain Subjects” to combine all results from two studies(43)).

Fear can also produce pain on time scales shorter than changes in sensitization. One form of fear that has been shown to heighten both pain responses and intensity is pain catastrophization, characterized by negative cognitive-emotional responses to actual or anticipated pain(30). Chronic pain and fear conditioning are intricately connected, suggesting that spinal pain-related circuits and cortical or subcortical fear circuits may undergo some of the same neural alterations during the transition from acute to chronic pain(34). Moreover, fear of pain is a consistent predictor of elevated pain intensity and greater perceived disability(49). Strategies that reduce pain catastrophization and strengthen a sense of safety can attenuate pain sensitivity and unpleasantness(50). Fear of pain may also be driven by anxiety, particularly in individuals with high anxiety sensitivity, a trait that amplifies fearful responses to potentially anxiety-inducing sensations(51). Anxiety itself can lead to increased perceived pain severity and decreased pain tolerance(52).

Neurons involved in the neural circuit attractor for chronic pain are present in many brain regions, and presumably throughout the entire pain processing system. The brain regions have been studied using fMRI by comparing blood-oxygen-level-dependent (BOLD) signals in control subjects and chronic pain patients(53,54). In individuals with chronic pain, researchers have observed hyperactivity within the default mode network (DMN), particularly in the medial prefrontal cortex (mPFC), as well as structures implicated in pain processing(2), including regions of the anterior cingulate cortex (ACC), nucleus accumbens (NAc), insula, and amygdala. It has been found that intrinsic network activity reflects the ongoing experience of chronic pain(55). This recent work complements and extends the pioneering work of Apkarian and colleagues, together with findings from other laboratories employing fMRI to identify alterations in brain activity that predict the transition from acute to chronic pain(20,56). Our simple model demonstrates that it is not the magnitude of signals from nociceptors or neuropathy alone that create the experience of chronic pain, but rather the altered processing of these signals due to the influence of sensitization, anxiety, and fear (SAF).

### 2.2. Development of Firing-Rate Model Parameters and Equations

We introduce two distinct time constants for Pain, *P*, and SAF, *S*: *τ*_*P*_ = 0.1 second(57) and *τ*_*S*_ = 1 week. Fortunately, the model is insensitive to the exact value of *τ*_*P*,_ which depends, of course, on the type of nerve conduction and the location of the pain stimulus on the body. Even changing *τ*_*P*_ to 10 seconds has a negligible effect on model predictions. The model is also relatively insensitive to the value of *τ*_*S*_, which is the time scale on which SAF can change. We chose 1 week as a compromise between the relatively rapid changes in Pain from changes in anxiety (Fig 2) and the slower changes in Pain from chronic pain recovery programs (typically 4 weeks or more). In future work, it may be worthwhile to enrich the model to separate these contributions, but we have chosen a low complexity, 2-dimensional, standard neural firing rate model in the interests of simplicity and parsimony. We normalize both Pain and SAF to values between 0 and 10; accordingly, we define the following sigmoidal activation function *f* (*x*) = 10/(1+*e*^*-x*^). As discussed above, the pain stimulus, ps, and the calming stimulus, cs, are input signals into the dynamics of pain and sensitivity. Overall, these two coupled nonlinear differential equations constitute a firing-rate recurrent neural network with excitatory connections from *P* to *S* and vice versa, and with an excitatory input ps and an inhibitory input cs.

Two main software programs were developed to (1) simulate the proposed dynamical system and (2) optimize its parameters. Standard Python libraries and routines were used. Specifically, we used the solve_ivp routine (with the LSODA numerical integrator) and the minimize routine (with the L-BFGS-B optimization algorithm). ChatGPT was used to automatically document code and design a few simple accessory routines. Three members of our team (FB, GM, TvdM) independently verified that the few accessory routines generated by ChatGPT are performing as intended.

For model predictions of chronic pain development, we extracted Apkarian’s data from figures in their paper (20) using the software WebPlotDigitizer and plotted it as described in the caption to Figure 1B. We used the supplemental equations *cs* = (10 – *cs*_*final*_)*e* ^-t/τc^ + *cs*_*final*_ and *ps* = *Ae* ^-t/τp^ to fit the calming stimulus, cs, and pain stimulus, ps. This gave us 5 parameters to fit:τ_*c*,_ τ_*p*,_ *cs*_*final*,_ *A* and *w* (see equation in 1B for *w*). We used an ODE solver (LSODA) from Python’s SciPy library to numerically integrate the equations and optimize the parameters to minimize the mean squared error from the target points of the Abkarian data. The result was:

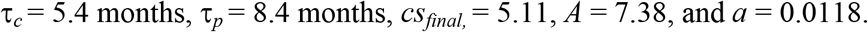

For model predictions of chronic pain recovery, we used data from the Boulder study(2) and the supplemental equation *cs* = (*cs*_*initial*_ – 10)*e* ^-t/τ’*c*^ + 10 to fit the parameters *cs*_*initial*_ and τ’_*c*_ . The result was *cs*_*initial*_ = 5.2 and τ’_*c*_ = 4.6 months. For development of chronic pain with different calming stimulus, we used the same values of τ_*c*_, τ_*p*,_ *w*, and *A*, but plotted the results for various values of *cs*_*final*_. For pain vs. stimulus, we simply used the Pain equation (Fig 1B) at steady state (dP/dt = 0) with values of SAF = 0 and 4.

For model predictions from our studies, we used data from our Study A and the same equations.

### 2.3. Participants and Enrollment

Participants with pain lasting longer than 6 months were recruited through clinician referrals, email lists, and posters. Inclusion criteria included a minimum age of eighteen, pain persisting longer than 6 months, and a minimum average pain level of 4 measured by the Visual Analog Scale(58). For Study A, recruitment began on 5/31/2023 and ended on 6/26/2023. For Study B, recruitment began on 3/22/2024 and ended on 4/19/2024. More details regarding participant enrollment can be found in the supporting information content flow chart (Fig S3). Participant demographics, including age, sex, race, and pain history, were collected (Table S2). Once participants consented to the study, they were assigned a weekly group meeting time. This study was approved by the Human Subjects Committee at UCSB (Study A protocol 3-22-0768 and Study B protocol 10-23-0730). Written consent was obtained for all participants.

### 2.4. Recovery Study Design

The study began with an initial individual meeting (in-person or Zoom) followed by weekly group Zoom meetings for the 4-week study duration. Each weekly Zoom meeting consisted of educational PowerPoint presentations with group discussions on the presented content. Participants were also assigned educational videos and worksheets to complete before each weekly Zoom meeting to reinforce their understanding of the material. Participants were provided with a hand-temperature biofeedback device for use at least twice daily. The study textbook was “The Way Out” by Alan Gordon in Study A and “The Explain Pain Handbook: Protectometer” by Lorimer Moseley in Study B. There were three online Qualtrics surveys: the initial application survey, at the end of the study, and 6 months post-study. These surveys assessed pain levels using the Visual Analog Scale(58) and Short-Form McGill Pain Questionnaire (SF-MPQ)(59) and assessed anxiety using the Short-Form State-Trait Anxiety Inventory (SF-STAI)(60).

### 2.5. Biofeedback Technology

The hand-temperature biofeedback device used in this study, the CalmStone, monitors changes in hand temperature. It was small enough to fit in the user’s palm for use in any setting. Its display comprised a ring of 6 lights that changed color one by one in a clockwise direction as hand temperature increased, indicating greater blood flow as a result of vasodilation that occurs during relaxation. It also had a breath pacer: the lights got brighter for breathing in and dimmer for breathing out. This device has not been approved by the FDA, and its use is still investigational.

### 2.6. Statistics and Reproducibility

To demonstrate that changes in anxiety were associated with changes in pain following biofeedback (in other words, that increased calming stimulus may predict decreases in the subjective experience of pain), we re-analyzed data from Studies 2 and 3 reported in Ly et al. (2023)(43), combining multiple biofeedback sessions from 15 participants across two 4-week studies (Fig 2). Here we fit a Bayesian hierarchical model with nested random effects (participants within studies) using brms(61) and Stan(62) in R. The technical minutiae of this modeling approach, including relevant prior specifications, have been described previously(43). However, in brief, we regressed the change in self-reported pain (before vs. after each biofeedback session, on a 0-10 VAS) against the change in anxiety (self-reported using the 6-item STAI). The nested random effects structure included both a random intercept and random slopes across participants/studies. Robust exploration of the posterior parameter space for all models was achieved using Hamiltonian Monte Carlo, with four independent chains each comprised of 15000 samples (the first 5000 of which were used for warm-up tuning of sampling parameters). For statistical inference, we report the population-level slope (given as the posterior median) along with the 95% highest-density interval around the median (Fig. 2).

We further sought to assess changes in pain observed across two new biofeedback studies (see **Study Design** above). Here we simply tested for differential reductions in pain between Studies A and B using an unequal-variance *t*-test (i.e., Welch’s *t*-test), given self-reported average pain levels before and after the study periods. We note that we also collected a more extensive series of self-report measures along with long-term follow-up data (6 months later). However, because not all participants were available to provide follow-up responses, we provide a descriptive visualization in Figure S2. Quantitative comparisons of these additional measures, before and immediately after each study, were obtained via mixed ANOVA (Table S1).

## 3. Results

### 3.1. Model Predictions

Our firing-rate model (Fig 1) is designed for simplicity with a small number of parameters determined by best fit to data from a landmark longitudinal study of chronic pain development conducted by Vania Apkarian’s group(20) (Fig 3A) and from the Boulder Study(2), a recent breakthrough investigation of chronic pain recovery by Tor Wager’s group (Fig 3B). The progression from acute to chronic pain depends on the magnitude of calming stimulus (Fig 3C). Even in the absence of pain stimulus, inadequate calming stimulus to counter sensitization, anxiety, and fear (SAF) can cause chronic pain to develop. Our model predicts that chronic pain can be reduced by supplying additional calming stimulus to diminish sensitization, anxiety, and fear (Fig 3B). The model provides a quantitative explanation of the interactions between Pain, SAF, pain stimulus, and calming stimulus. More details on the development of the model can be found in Methods.

**Fig 3.**
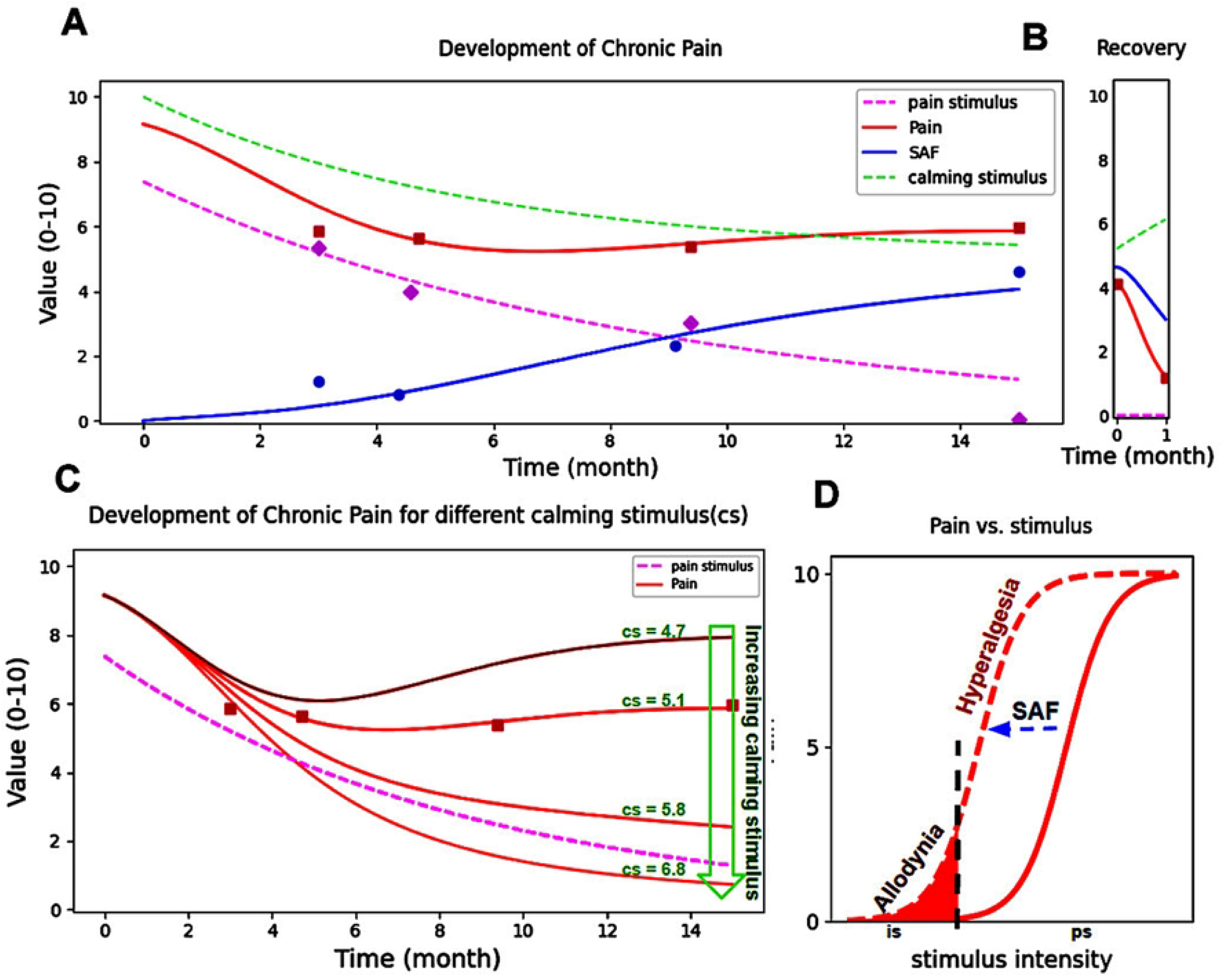
Model results from the equations in the firing-rate recurrent neural network. **A**. Parameter fitting based on neuroimaging data from Apkarian’s group(20). “Pain” data comes from their Figure 3B. “Pain stimulus” data comes from their Figure 3D, based on their mask for acute pain in which SAF is zero. “SAF” data comes from their Figure 3D, based on their mask for emotion, which increases as chronic pain develops, and is used here as a measure of SAF. Equations and methods for fitting the calming stimulus and pain stimulus are detailed in Methods. **B**. Data for fitting recovery comes from the Boulder Study(2). An exponential increase in calming stimulus was used due to the absence of intermediate data points to fit. **C**. In the model, the development of chronic pain is strongly dependent on calming stimulus: greater calming stimulus results in decreased pain. This emphasizes the need for calming stimulus as part of rehabilitation programs. **D**. The first equation of the model reproduces the well-established dependence of Pain on pain stimulus. The zero for pain stimulus, ps, is set at the onset of pain. Less intense stimulus is labeled “is” for innocuous stimulus. More details can be found in Methods.

Model predictions using these parameters were consistent with the human study results reported in Figure 4A for a greater variety of chronic pain conditions. A study narrowly focused on reducing pain-related anxiety and fear produced an average pain decrease of 61% or 3.5 (on a 0 to 10 scale), with half the subjects who adhered to the recovery program ending up with little or no pain (1 or 0) (Fig 4A).

**Fig 4.**
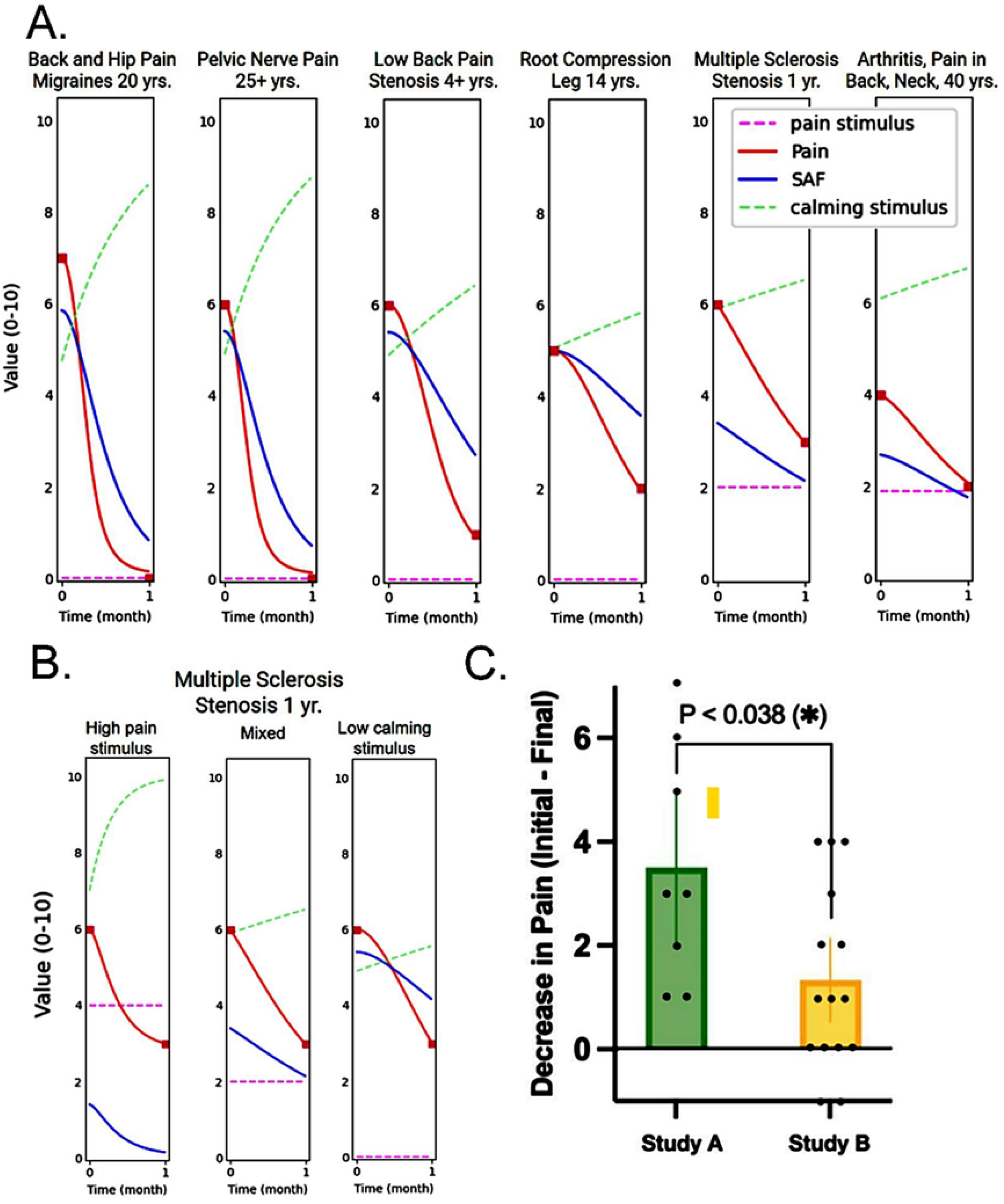
Recovery for various chronic pain conditions based on the model equations. **A**. Model fits for the six subjects who did the assigned work in Study A. The two on the left can be fit with some confidence since they ended up with no pain, which tells us they had no pain stimulus. For the remaining four, it is unclear whether they ended up with some pain due to pain stimulus, inadequate calming stimulus, or a combination of both. Since people with low back pain, stenosis, and root compression can sometimes end up with no pain(2), we assumed, for the graphs, that their problem was inadequate calming stimulus. Since arthritis and multiple sclerosis could be expected to result in pain stimulus, we assumed a non-zero pain stimulus. The labels are summaries of the data the subjects supplied in their applications (complete data in Fig S1.). **B**. Three different possibilities for the subject with multiple sclerosis: a high pain stimulus with adequate calming stimulus (left), no pain stimulus, but inadequate calming stimulus (right), or “Mixed” with both some pain stimulus and inadequate calming stimulus (middle). Note that the curvature of the Pain curve is different for these different possibilities. If calming stimulus is adequate, the pain decreases rapidly at first (left). If calming stimulus is inadequate, pain decreases more slowly at first (right). **C**. Average Pain, defined as the average self-reported pain in the previous week, decreased from study application (Initial) to study conclusion (Final). The decrease was nearly twice as large for Study A (n=8) compared to Study B (n=15)(*g* = 1, 95CI = [0.06, 1.90]). Error bars give 95% confidence intervals around the mean decrease for each study.

### 3.2 A Comparison of Two Strategies for Increasing Calming Stimulus

In the model, the key to reducing nociplastic pain is increasing calming stimulus, which reduces pain-related sensitization, anxiety, and fear. We compared two different strategies for increasing calming stimulus by conducting two preliminary pilot studies (A and B). Figure 4C shows that pilot study A resulted in a greater decrease in pain than study B. The studies differed in their use of educational textbooks ( *The Way Out* versus *The Explain Pain Handbook: Protectometer*), participant numbers, and the inclusion of Zoom breakout rooms in Study B. Additional smaller studies aimed at isolating the influence of these variables would be valuable to clarify their respective contributions.

## Discussion

Our simplified model explains how chronic pain can occur even in the absence of tissue injury that would cause a pain stimulus. It further explains how chronic pain can be effectively addressed with non-pharmacological interventions grounded in the fundamental neurobiological principles presented here–specifically, by reducing the fear and anxiety associated with sensation. The model makes specific predictions such as the presence and time dependence of an fMRI signal related to calming stimulus. It is a falsifiable model: future studies could confirm or refute its predictions and refine its applicability. It may help with the design of future neural circuit interventions (63).

The results observed in our previous chronic pain recovery studies(43) and in the two new studies reported here support our proposed model for chronic pain. Overall pain was greatly reduced by attenuating the sensitization, anxiety, and fear involved in chronic pain when calming stimulus (cs) was supplied for many different types of chronic pain (Fig 4A). Although the specific neural circuits and mechanisms underlying chronic pain are not completely understood, evidence from our work and other studies(2) demonstrates that an understanding of the causative factors in chronic pain is already sufficient to design a program to reduce overall pain. While cellular pathophysiology is crucial for pharmacological interventions, extensive research has shown that pain medications are largely ineffective for chronic pain due to neuroplastic changes. Therefore, treatment strategies can shift toward understanding chronic pain mechanisms at a therapeutic level, as outlined in our model. Detailed knowledge of the specific neural circuits, biochemical mechanisms, and neurotransmitters involved in pain due to neuroplastic pain may be helpful, but is not required to alleviate pain-related anxiety and fear, which is the key to reducing pain.

Results from our study, reported here, along with other studies, suggest that most individuals experience reductions in chronic pain when they experience reductions in pain-related fear and anxiety, regardless of the type of chronic pain. This indicates that extensive diagnostic procedures for nociplastic pain may not always be necessary, particularly when they create barriers to initiating chronic pain recovery, such as limited access to specialized physician training or vetted tests. For example, Clauw has emphasized that Quantitative Sensory Testing (QST) has significant limitations: 30% to 70% of patients with fibromyalgia, a prototypical nociplastic pain condition, lack evidence of central sensitization on QST(64). Recognizing that nociplastic mechanisms are likely present in the majority of chronic pain cases, including chronic pancreatitis(65), could allow clinicians to more readily direct patients to effective treatment programs. Such an approach has the potential to substantially reduce the billions spent annually on prescription pain medications while mitigating the risks of misuse and addiction associated with long-term opioid therapy(66).

This model points to an important research opportunity: exploring the most effective methods for targeting anxiety and fear associated with sensation. A progressive research approach employing targeted, comparative studies of individual program components will help identify the most effective methods. Such advances could improve validated, resource-worthy programs suitable for physician referral and insurance reimbursement.

## Acknowledgements

We thank Dahyana Arroyo for acting as a study coordinator for Studies A and B. We acknowledge support from John and Kathy Kirtley. TS and MBM were supported by the Army Research Office under contract W911NF-19-D-0001 for the Institute for Collaborative Biotechnologies. We thank Ericka Dickson, Hal Kopeikin, and Pamela Benham for useful suggestions.

None of the authors have any financial connection with the company that makes the CalmStone biofeedback devices or the textbooks used in this study. The devices for this study and the textbooks were purchased, not donated.

All data are available upon reasonable request to the corresponding author (PH).

## Code Availability

The source code is available at https://github.com/GuyMarmor/ChronicPainModel. The GitHub repository contains Python code for modeling the development of and recovery from chronic pain fitting data from the Apkarian group’s fMRI studies, the Boulder Recovery Study, and our studies (A and B). The source code is licensed under an Apache License 2.0 and can be found using the DOI 10.5281/zenodo.3482768.

## Supplemental Digital Content Figure and Table Captions

**Fig S1. Conditions, pain duration, and change in pain for Studies A and B**. Descriptions of type, location, and duration were copied verbatim from participants’ application forms. In both studies, most participants experienced decreases in average pain. **a**. Data for Study A participants. **b**. Data for Study B participants.

**Fig S2. Figure S2. Self-reported pain, anxiety, and recovery metrics at three time points (baseline, post-intervention, and 6-month follow-up) comparing Study A and Study B**.

**a**. The Visual Analog Scale (VAS) Minimum Pain captures the self-reported minimum pain during a typical week on a 0 to 10 scale, comparing results between Study A study B. All initial scores represent the minimum pain level upon applying to the study, the 1-month scores represent the minimum pain level after completing the 5-week study, and the 6-month scores represent the minimum pain level at the 6-month follow-up. **b**. The Visual Analog Scale (VAS) Average Pain captures the self-reported average pain during a typical week on a 0 to 10. **c**. The Visual Analog Scale (VAS) Maximum Pain metric captures self-reported maximum pain during a typical week on a 0 to 10 scale. **d**. The Visual Analog Scale (VAS) Current Pain metric captures self-reported current pain intensity on a 0 to 10 scale. **e**. The Short Form McGill Pain Questionnaire (SF-MPQ) captures the degree or severity of the participant’s pain by summing the self-reported ratings from 0 (none) to 3 (severe) for each of 15 descriptive adjectives (throbbing, shooting, stabbing, etc.). **f**. The McGill Present Pain Index (PPI) captures self-reported current pain intensity on a six-point scale from 0 (no pain) to 5 (excruciating). **g**. The Satisfaction and Recovery Index (SRI) metric captures self-reported recovery from participants’ ailments in terms of interference from injury and symptoms, generating a percent health-related satisfaction score. Higher percentages represent higher satisfaction. **h**. The Short-Form State-Trait Anxiety Inventory for Adults (STAI-AD) captures levels of current state anxiety by asking participants to select between “not at all” and “very much so” for 10 descriptive emotional statements in the present moment. A higher value represents a higher level of anxiety.

**Table S1. Change in average pain levels before and after the biofeedback program, separated by study**.

In Studies A and B, most participants, even those with underlying disease, experienced decreases in average pain (Fig S1). The studies focused on decreasing *sensitization-anxiety-fear* with calming stimulus. The six subjects above the dotted gray line in Study A did the assigned work and had a greater decrease in pain. Compared to Study A, all Study B participants did the assigned work, but had a lower reduction in pain, possibly due to the addition of increased pain exposure via an activity goal or less effective calming stimulus.

**Table S2. Mixed ANOVA results comparing changes in pain, anxiety, and recovery metrics between Study A and Study B before and after the intervention**.

A mixed ANOVA test revealed a statistically significant difference in 4 out of 8 total metrics, 6 of which measure pain, when comparing change in initial scores and final scores (completion of 5-week study) between studies A and B. The statistically significant metrics included average pain, minimum pain, current pain, and the Short-Form McGill Pain Questionnaire (SF-MPQ). No significant differences were found when comparing the change in maximum pain, Present Pain Index (PPI), State-Trait Anxiety Inventory for Adults (STAI-AD), and Satisfaction and Recovery Index (SRI). The latter two are not pain-related metrics. Partial eta-squared is a reliable measure of effect size. Though context-dependent, commonly accepted benchmarks for small, medium, and large effect sizes are approximately 0.01, 0.06, and 0.14, respectively.

**Table S3. Demographic and clinical characteristics of participants in Study A and Study B**.

Demographics and Clinical Characteristics are separated by study.

**Fig S3. Flow diagrams for participant enrollment in Studies A and B**.

**a**. Flow Diagram for Participant Enrollment for Study A. In addition to subjects with chronic pain, one individual with self-reported persistent “burning dry mouth and terrible oral salty taste” after administration of the COVID-19 booster was enrolled on a trial basis. Since he did not complete all required assignments and his symptoms did not alleviate, he was not included in this flowchart or the data analysis. **b**. Flow Diagram for Participant Enrollment for Study B. Initial Inquiry: submission of preliminary Google Form to determine study eligibility. Did not complete study: did not attend the minimum required number of Zoom education sessions or did not complete the minimum required number of biofeedback sessions or did not submit the minimum required number of Weekly Worksheets.

